# Extensive splicing deficiency in a degenerating mating-type chromosome

**DOI:** 10.64898/2025.12.09.693058

**Authors:** Chris Condon, Andrea Galvez, Alexander Kramer, Landen Gozashti, Chris Vollmers, Manuel Ares, Russell Corbett-Detig

## Abstract

Splicing deficiency may represent a critical yet underexplored form of genomic erosion in non-recombining regions. Across four phytoplankton species diverged ∼333–639 million years ago, genes within U (female) and V (male) “UV” mating-type regions—non-recombining chromosomal regions that determine mating compatibility—show strikingly elevated intron retention relative to genes in other genomic regions. Long-read data reveal abundant aberrant, likely non-functional mRNA isoforms despite preserved coding potential. This preservation suggests that splicing defects arose early in UV evolution and have persisted over deep time. We propose that these defects arise from evolutionary changes in sequence composition and chromatin organization that accompany recombination suppression, such as reduced GC content, altered nucleosome occupancy, and disrupted methylation, that collectively compromise splicing fidelity. Unlike sex chromosomes, which often degenerate through gene loss, splicing-deficient UV regions in green algae retain hundreds of genes, indicating that transcript-level dysfunction provides an alternative route to functional decay. Our results identify chromatin-mediated splicing deficiency as a novel axis of genomic erosion and position algal UV systems as models for studying how recombination suppression reshapes RNA processing fidelity in essential, non-recombining genomes.

## Introduction

In eukaryotes, suppressed recombination drives predictable genomic decay including disrupted gene order, repetitive element expansion, and biased nucleotide composition [1,2]. Many studies focus on highly degenerate regions like mammalian Y chromosomes, where extensive gene loss reflects relaxed selection on non-essential genes (*e*.*g*., [3]). However, many non-recombining regions must retain essential functions—such as mating-type loci in fungi and algae, centromeric regions, and loci harboring large segregating structural rearrangements—creating evolutionary tension between genomic decay processes and strong functional constraints [4,5]. Understanding how this tension resolves has broad implications for genome evolution and for identifying the factors that drive the emergence of genetic innovation.

UV mating-type (MT) regions in the green algae offer a compelling model for studying genomic erosion with selective constraints to maintain genetic function. These largely non-recombining regions span 393-1,681 kb across the four Mamiellales species examined here, and contain 216-619 genes [6]. The boundaries of these regions are defined by sharp transitions in GC content, which is a hallmark molecular signature of recombination suppression resulting in the loss of GC-biased gene conversion [6]. Despite sharing 23 core gene families that define orthologous UV loci across Mamiellales, these regions show substantial species-specific gene content and structural divergence. Within O. tauri, the MT- and MT+ alleles differ in size (∼650kb vs. ∼450 kb) and contain both mating-type-specific genes and 75-79 shared genes beyond the core gene families.

Like non-recombining sex chromosomes, UV loci show disrupted synteny, lower gene density, expanded repeat content, decreased GC nucleotide content [5–7], and more open chromatin [8] compared to autosomes. Notably, UV loci in *Mamiellales* also show higher gene expression than the autosomal regions [7,8]. Comparative genomic analysis provides evidence for past translocations in O. tauri, where 46 gene families in MT- and 30 in MT+ have paralogs located on diverse autosomes [6]. Unlike the usually highly degenerate Y or W chromosomes, UV loci encode hundreds of actively expressed genes with critical biological functions [7], suggesting that degeneration is tempered by selective pressures and may manifest in fundamentally different ways from more commonly studied sex chromosomes.

We hypothesize that transcript processing dysfunction represents a cryptic form of erosion in functionally constrained regions. RNA splicing fidelity depends on sequence context including both consensus splice sites and auxiliary splicing enhancers and silencers [9–11], chromatin structure [12], and DNA methylation patterns [13,14]. Recent comparative studies demonstrate that species with smaller effective population sizes accumulate more rare, likely non-functional alternatively spliced transcripts, suggesting that reduced efficacy of natural selection permits aberrant splicing [10]. If suppressed recombination similarly degrades splicing while preserving coding sequences, this could diminish gene function without complete gene loss—a form of “molecular erosion” largely invisible to traditional genomic analyses.

Using comparative transcriptomics across four Mamiellales species, we find widespread splicing dysfunction within recombination-suppressed mating-type regions. Long-read isoform sequencing further demonstrates that this dysfunction produces an excess of aberrant, likely non-functional transcripts, providing evidence for transcript-level genomic erosion. Our results reveal transcript-level defects as a novel axis of genomic decay and establish the mating-type regions of Mamiellales as a model for studying transcriptional dysregulation.

## Results

### Splicing In Green Algae Mating-Type Region is Highly Aberrant

Transcriptome analysis among four Mamiellales species revealed widespread and apparently conserved splicing misregulation within the mating-type region (Figure 1). We compared splicing patterns between previously identified mating-type regions to other regions in four Mamiellales species [6]. In *Micromonas pusilla* the mean intron retention frequency outside of the mating-type region was 0.011, while within the mating-type region the mean frequency was 0.33 (p < 0.0001, Mann-Whitney U). We estimate that nearly half (45%) of all transcribed introns derived from genes within the mating-type region are retained in these data. This difference (33% vs 45%) indicates that the frequency of intron retention is more common in transcripts of highly expressed genes (see also below). Despite approximately 339-630 million years since the last common ancestor of these taxa [15], this pattern is qualitatively consistent across each species (Figure 1, Table S1), implying that the processes that contribute to this apparent aberrant splicing have been operating since shortly after the emergence of the UV mating-type region and have been retained in these diverging lineages.

**Figure 1.**
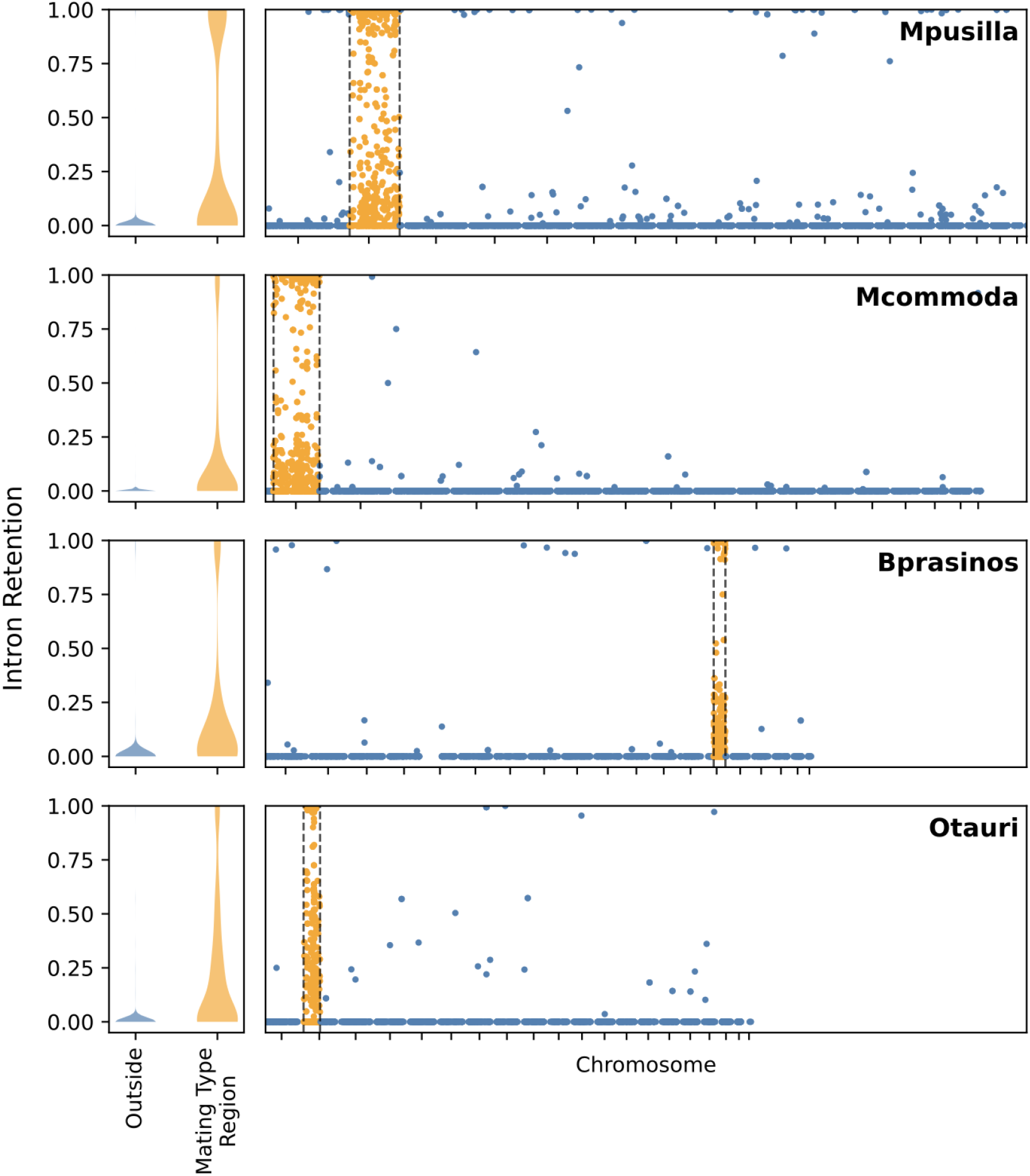
Individual intron retention patterns across the genome of Mamiellales species. (Left), per intron retention frequency along autosomal regions (blue) and within the mating-type region (orange). (Right) Retention frequency for each intron plotted along each chromosome with colors as on the left panel. We obtained coordinates for mating type regions from [6]. Tick-marks indicate the midpoints of chromosomes in each genome assembly and we excluded any contig below 50Kbp.

### Possible Mechanistic Basis of Aberrant Splicing

Introns in mating-type genes are highly divergent from those elsewhere in the genome. They are significantly shorter with lower GC content (Figure S1), both features that affect splicing efficiency [16,17]. Most strikingly, branchpoint sequences which are critical for spliceosome assembly, are significantly depleted in all species within the mating-type region (p < 0.0001 for all species; Table S3). Splice-site flanking sequences also reflect the broader GC depletion in the mating-type region, though specific positions within the extended splice site consensus motifs retain their nucleotide compositions (Figure 2, Figure S2). Splice-site flanking sequences reflect the broader GC depletion in the mating-type region, though specific positions within the extended splice site consensus motifs retain their initial nucleotide compositions (Figure 2, Figure S2). For example, guanine at the fifth intron position of the 5’ splice site—which contributes to splicing efficiency by base pairing to U6 snRNA [18–20]—is significantly less frequent in mating-type introns (8%-23% reduction; *p* < 0.0001 for all species, Fisher’s exact test; Table S2), yet these sites remain enriched for guanine relative to other intronic positions, suggesting that natural selection is relaxed but detectably present. In contrast, canonical splice sites (5’ GT/GC and 3’ AG) occur at similar frequencies inside versus outside the mating-type region (Figure S2), with only two subtle exceptions: *M. commoda* shows slightly fewer canonical 3’ splice sites in mating-type genes (96.1% vs. 98.6%, p < 0.0001), while *M. pusilla* shows slightly more canonical 5’ splice sites (96.6% vs. 98.2%, p=0.41E=3). These results indicate that while mating-type introns differ markedly in composition and length, natural selection maintains core splicing signals but acts less efficiently on secondary elements that affect splicing efficiency.

**Figure 2.**
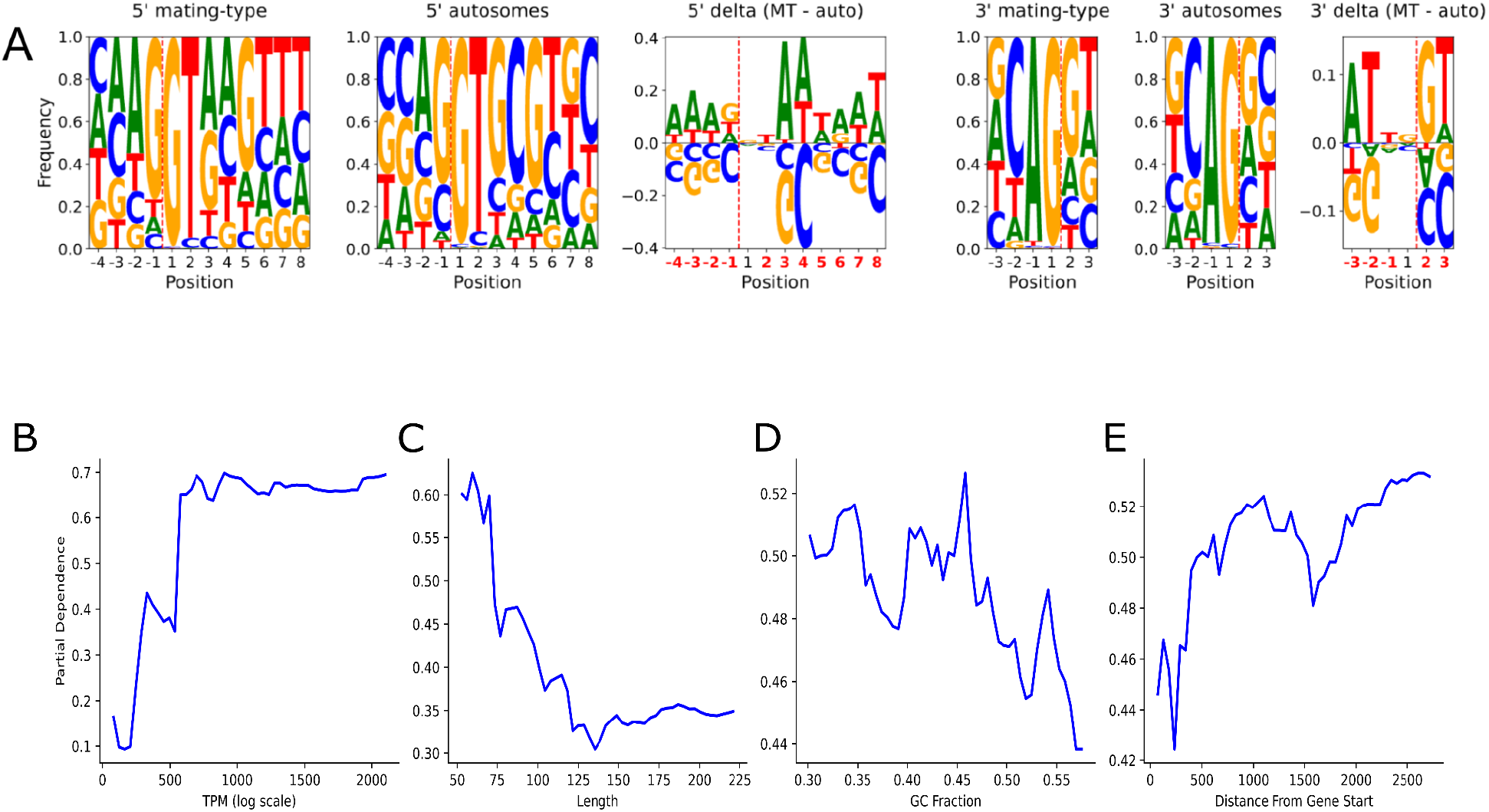
Splicing-related features for *M. pusilla*. (A) 5’ and 3’ splice sites in the mating-type region, 5’ and 3’ splice-sites in autosomal regions, and the difference in sequence composition between mating-type and autosomal regions’. Intron boundaries are denoted with a dashed vertical line. Partial dependence plots for (B) TPM, (C) intron length, (D) GC fraction, and (E) intron distance from the gene start. The Y-axis shows the marginal effect of each feature on predicted intron retention probability, holding all other features at their average values. Higher values indicate increased likelihood of retention. Feature importances are provided in Table S4.

Random forest modeling considering only introns within the mating-type loci confirmed that shared transcript and genomic features predict the occurrence of splicing dysfunction across all four species (AUC-ROC: 0.735–0.867). Transcripts per million, a measure of gene expression, was the strongest predictor, wherein highly expressed genes showed the highest intron retention rates across all species. Shorter introns, lower GC content, and greater distance between the intron and the transcription start sites also predicted retention (Figure 2, Table S4), while potential functional consequences (frameshifts, stop codons) and the presence of specific splicing features (branchpoints and splice-site adjacent nucleotides) showed little predictive power and were rarely retained in final models after applying minimal feature selection techniques. In contrast, Random Forest models trained on autosomal introns failed to achieve meaningful predictive performance (F1=0.00-0.17) due to extreme class imbalance where retained introns comprise only 0.6-2.6% of autosomal introns (Table S4). The absence of predictive power for frameshifts or stop codons may reflect active nonsense-mediated decay (NMD). Core NMD machinery (UPF1, UPF2)[21] is expressed across all species (Table S7) and transcripts with retention induced frameshifts or stop codons would be primary degredation targets. Our measurements therefore likely capture the subset of retained introns that saturate or escape NMD. Notably, 16–35% of variance in intron retention occurred at the gene level across all species, indicating coordinated splicing regulation within genes rather than independent intron-by-intron effects. Consistent with this coordination, intron number per gene was strongly correlated with retention rate in mating-type genes (Spearman’s ρ = 0.56–0.65, p < 0.0001) but not in autosomal genes (ρ = 0.06–0.22), suggesting that splicing defects compound across multiple introns specifically in the mating-type region.

The mating-type region of *M. pusilla* is substantially different from the other genomic regions with respect to the local chromatin environment [8]. MNase-based measurement of chromatin accessibility [22] suggests that genes residing in the mating-type region have substantially altered nucleosome occupancy. Relatedly, there are fewer CpG dinucleotide sites in exonic regions flanking each intron within the mating-type region consistent with lower overall GC nucleotide content (Figure S3). Nonetheless, CG dinucleotides that are present within the mating-type region are significantly less likely to be methylated on a per-site level than are those in the autosomal regions of the genome (Figure S3). This implies that overall GC content and methylation efficiency are critical determinants of differences in methylation patterns.

### Diverse Isoforms are Created by Widespread Missplicing

Analysis of full length isoforms in *M. pusilla* produced detailed insights into mRNA processing at the transcript level. We circumvented the vagaries of interpreting short-read RNA sequencing data by applying a long-read circularized consensus method [23] to generate 14,350 high quality full length isoforms for the *M. pusilla* reference strain. Using a linear regression model, isoform number increases with both expression and intron count (0.27 isoforms per log-expression unit, 0.76 per intron). Genes in the mating-type region produce ∼1.9 more isoforms than autosomal genes, with predicted means of 1.83 (non-mating-type region genes) vs. 3.72 isoforms for mating type region genes at average expression and intron number (*p* < 0.0001, OLS; Figure 3). Additionally, genes within the mating-type region show more evenly distributed isoform expression, with less dominance by a single major isoform, compared to autosomal genes (p < 0.0001, GLM; see Methods). This pattern suggests that purifying selection is less efficient at suppressing non-canonical isoforms in the mating-type region.

**Figure 3.**
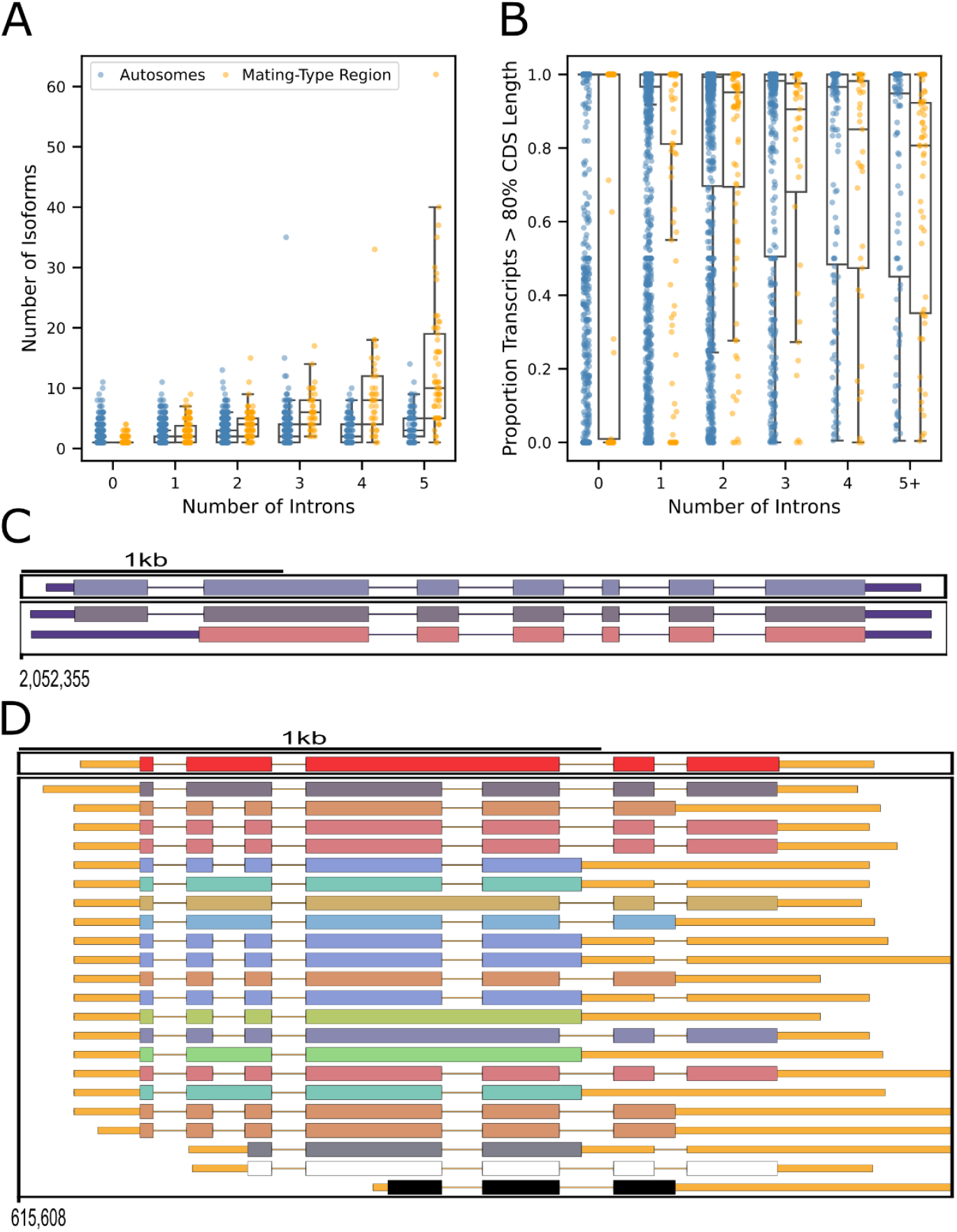
Isoform diversity within and outside of the mating-type region in *M. pusilla*. (A) The number of isoforms per gene in mating-type (orange) and autosomal genes (blue). (B) The proportion of transcripts that contain a minimum of 80% of the CDS of the canonical gene’s CDS. In A and B all genes with more than 5 introns are collapsed and displayed as 5+ introns.(C) The complete isoform set from a representative gene, MicpuC2.PR_0200000013, selected from the autosomal regions. (D) The complete isoform set from a representative gene, MicpuC2.EuGene.0000020604, selected from the mating-type regions. Each horizontal track represents a distinct isoform (colors assigned for visual distinction). Thick colored bars indicate exons, thin black lines indicate spliced introns, and orange medium-width lines indicate untranslated regions (UTRs). Representative genes were selected based on the median number of isoforms for each region and genes with 6 introns.”

We also compared other features of transcript diversity including transcript start sites and polyadenylation sites for genes within and outside of the mating-type regions. This analysis revealed selective disruption of 3′ end regulation whereby genes in the mating-type region exhibit significantly more distinct transcript termination sites compared to genes outside the region (39.4% increase; *p* < 0.0001, Poisson GLM). This indicates that like splicing, termination and polyadenylation may be similarly disrupted. However, genes within and outside the mating-type region display comparable diversity in transcription start sites across isoforms (*p* = 0.563, GLM). We speculate that while the accumulation during evolution of AT-rich sequences may contribute to increased alternative polyadenylation, mispriming of oligo-dT primers during library construction could artifactually contribute to this pattern. Together, these findings suggest that co-transcriptional and post-transcriptional RNA processing—but not transcription initiation—are perturbed in the recombination-suppressed mating-type region.

Transcripts derived from genes within the mating-type region are less likely to be functional than those in other regions of the genome. In the absence of direct experimental validation, it is challenging to determine if a given transcript produces a functional protein. We therefore indirectly estimated functionality by asking what fraction of full-length transcripts are derived from isoforms that contain at least 80% of the annotated CDS in the correct reading frame without premature termination. Strikingly, transcripts from genes within the mating-type region showed a significantly lower fraction meeting this threshold (*p* < 0.0001, GLM, Figure 3B). To further assess the effect of alternative isoforms on protein function, we annotated functional protein domains and ontologies for each isoform. Isoforms derived from genes in the mating-type region also more frequently exhibit functional domain disruptions and changes in functional ontology compared to genes in the rest of the genome (Table S5). For example, for genes in the mating-type region, ∼87% of isoforms contained at least 90% of annotated functional domains on average, compared to >92% for genes in autosomal regions (*p* < 0.001, Mann-Whitney U; Table S5). These patterns suggest that the elevated alternative splicing observed in the mating-type region is primarily deleterious, though we cannot exclude the possibility that some alternatively-spliced isoforms are neutral or positively selected.

## Discussion

We propose that recombination suppression triggered a cascade of interconnected molecular changes and distinct selective pressures that collectively explain these widespread transcript-level defects (Figure 4). The suppression of recombination in the mating-type region dismantles the primary evolutionary mechanism that maintains genomic integrity against specific, underlying mutational pressures that have been empirically demonstrated in Mamiellales [24] and common to most eukaryotes [25]. This mutational pressure driving GC to AT transitions is counteracted in the recombining portions of the genome by GC-biased gene conversion (gBGC). The reduction of recombination in the mating-type region reduces gBGC, diminishing the main impediment. The strong intrinsic bias for GC→AT mutations drives the dramatic shift toward an AT-rich sequence composition. Simultaneously, a deletion bias causes a net loss of base pairs per generation. This effect systematically shortens non-coding regions, and due to decreased efficacy of selection in non-recombining genomics segments, introns gradually evolve to be shorter. The collective action of mutational factors and natural selection thereby result in a distinct nucleotide composition within the mating-type region.

**Figure 4.**
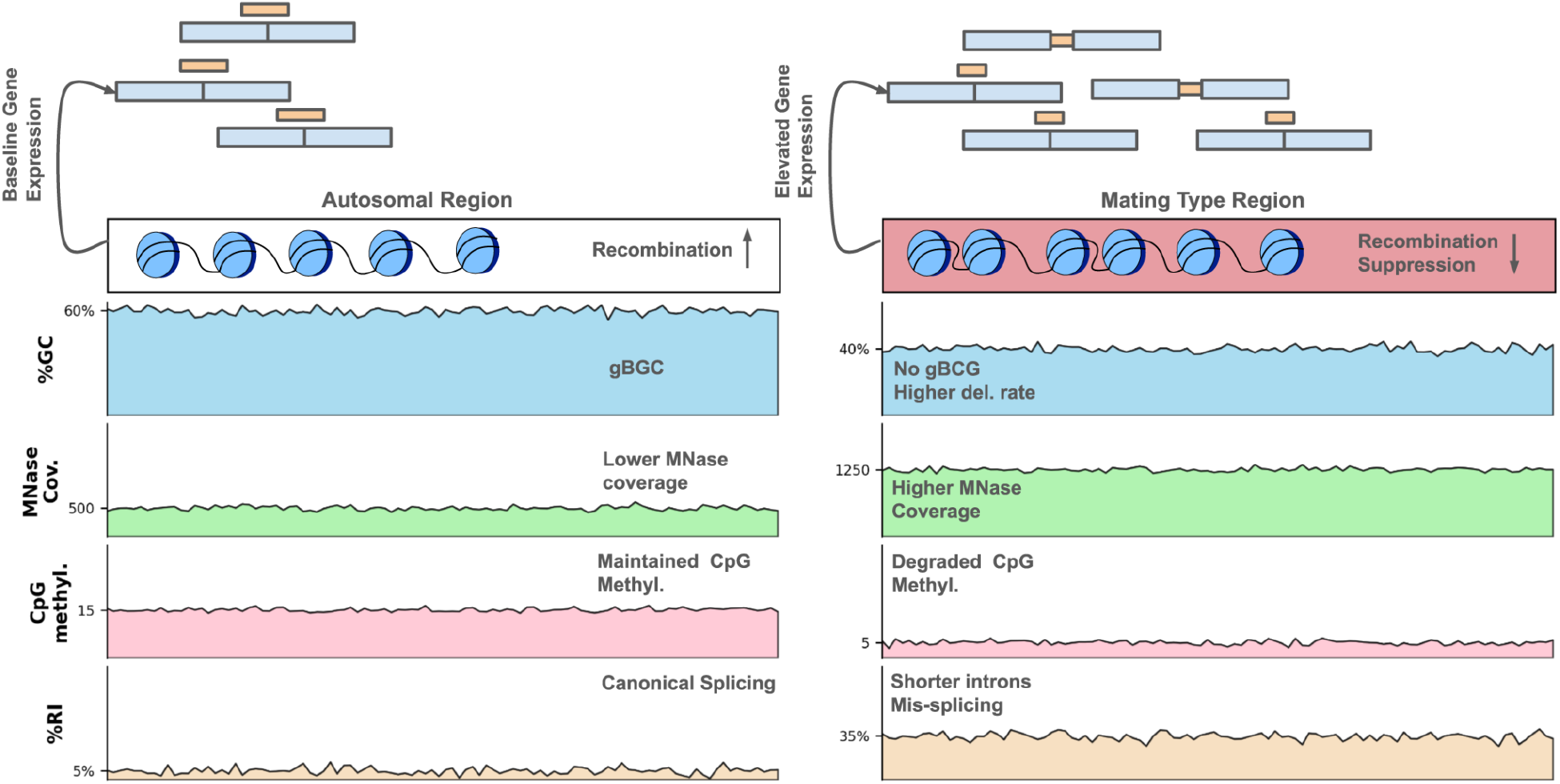
Proposed model of genomic erosion in the *Mammiellales* mating-type region. Schematic comparing autosomal regions (left) with mating-type regions (right), illustrating how recombination suppression initiates a cascade of molecular changes. Suppression of recombination eliminates GC-biased gene conversion (gBGC), allowing GC→AT mutational bias to reduce GC content from ∼60% to ∼40%. This compositional shift reduces CpG dinucleotide availability and methylation efficiency, disrupts nucleosome positioning (elevated MNase coverage indicates more open chromatin), and shortens introns through deletion bias. These chromatin and sequence changes collectively impair splicing fidelity: ∼35% of mating-type introns are retained (thick orange bars in gene models) compared to ∼3% in autosomal regions, producing abundant truncated, dysfunctional transcripts despite preservation of coding sequences. Expression levels (number of gene boxes) are elevated in the mating-type region due to altered chromatin accessibility, further exacerbating splicing defects through kinetic mismatches between transcription and co-transcriptional splicing.

We propose that this collective action of mutational biases and natural selection then causes decreased efficacy of splicing in the evolving mating-type region through a series of molecular consequences. First, this dramatic shift in GC content has compounding effects on chromatin structure: fewer CpG dinucleotides provide fewer sites for DNA methylation, while AT-rich sequences alter nucleosome positioning and consistency in the mating-type region [8,14,22,26]. Second, changes in GC content and nucleosome positioning can disrupt transcription kinetics essential for splicing efficiency and fidelity [27,28]. Third, altered chromatin structure appears to elevate overall gene expression levels in humans [12], consistent with our finding that highly expressed genes in the mating-type region show the highest intron retention rates across species. This positive correlation between expression and retention is paradoxical given that some species show the opposite pattern [10,29], where selection minimizes splicing errors in highly expressed genes. However, in the *Mammiellales* species considered here, gene expression and intron retention are weakly positively correlated even in autosomal regions, where selection can operate efficiently to minimize intron retention (Spearman’s ρ = 0.06-0.12 across species, p <0.0001). This suggests fundamentally different mechanisms may govern transcription and splicing fidelity across eukaryotes. Furthermore, in the mating-type region, altered chromatin structure enables a loss of kinetic coupling between transcription and splicing. More rapid transcription by RNA polymerase II through more altered chromatin might outpace or disrupt co-transcriptional spliceosome assembly, leading to inappropriate exon inclusion or skipping (e. g. [30,31]). Fourth, functional coupling between splicing and 3’ end formation can lead to transcript 3’ end heterogeneity when splicing is inefficient [32], an effect we also observe in transcripts from the mating type regions. This cascade provides a mechanistic basis for how non-recombining but functionally essential genomic regions accumulate progressive transcript-level defects while retaining the capacity to produce a fraction of functional transcripts sufficient to maintain viability.

A critical question remains why genes whose functions are unrelated to mating compatibility remain inside the UV region despite apparent persistent dysfunction. While comparative genomic analysis demonstrates that gene trafficking between UV and autosomal regions has occurred during Mamiellales evolution [6], we hypothesize that relocation alone does not immediately restore splicing function. In the event of a translocation, a gene would carry the sequence-level changes accumulated during its residence in the UV region, including degraded GC content that has been shown to persist in genes relocated from brown algal SDRs to autosomes [33]. Studies of X chromosome gene trafficking demonstrate that relocated genes require substantial evolutionary time to re-establish proper regulatory contexts [34,35]. Furthermore, genes would need to evolve compensatory mutations that restore lost splice site strength and intron length. This evolutionary lag may exceed the timescale over which selection maintains even partial gene function in situ, explaining why the system tolerates ongoing transcript-level dysfunction rather than rapidly relocating non-mating genes.

This mode of genomic erosion represents a fundamentally distinct evolutionary trajectory than the gene-loss paradigm exemplified by highly degenerate mammalian Y chromosomes. Rather than wholesale elimination of dispensable genes under relaxed selection, mating-type regions experience a form of molecular compromise: genes are retained but function at reduced efficiency due to pervasive splicing deficiency. The persistence of this pattern across more than 300 million years of Mamiellales divergence suggests that producing even a modest fraction of correctly spliced transcripts may satisfy selective constraints when complete gene loss would be lethal. This framework has broad implications beyond green algal UV systems. Other non-recombining but essential genomic regions, including the young sex chromosomes of many plant and animal species, large structural polymorphisms segregating in natural populations, and even highly repetitive centromeric regions, may similarly accumulate cryptic transcript-level dysfunction that precedes or accompanies more visible forms of degeneration. Our model generates testable predictions: recently formed sex chromosomes should show intermediate splicing deficiency that scales with the time since recombination suppression, and experimental restoration of recombination should improve splicing fidelity. These findings therefore establish Mamiellales mating-type chromosomes as a powerful comparative system for dissecting how genome architecture, chromatin state, and RNA processing fidelity interact in shaping the evolution of essential non-recombining regions across eukaryotes.

## Methods

### Software Development

We used LLM support to develop initial scripts and analyses. Each script was tested and ultimately validated by the authors prior to inclusion in this manuscript.

### Comparative Intron Retention Analyses

We identified publicly available short-read RNAseq data for four species in the Mamiellales family through metadata search on the sequence read archive (Table S1). We further restricted the search to species whose genomes are chromosome level assemblies with a mating-type region defined in [6]. We aligned RNAseq data to each genome using STAR [36] version 2.7.10b and provided the appropriate gtf file during index construction, and the “--quantMode GeneCounts” option to obtain estimates of gene expression, but used otherwise default parameters. We used rMATs [37] version 4.3.0 to quantify intron retention providing the parameters “--allow-clipping --novelSS” but otherwise default in running this analysis.

We fit a linear mixed-effects model of intron retention ratios, using gene ID as a random effect, to estimate the variance in intron retention explained by differences among genes and to calculate the intra-gene correlation.

### Branchpoint-Sequence Detection

We extracted introns from annotated CDS features in GTF files and retrieved the corresponding 50-nt upstream branchpoint windows from the reference genome, outputting unique sequences in FASTA format for downstream analyses. These sequences were then analyzed with DREME [38] to identify enriched short motifs. In each case, the most significantly enriched motif is consistent with expected branchpoint sequences in these species.

### Methylation and Chromatin Accessibility Data

For *M. pusilla*, we complimented gene expression estimates and intron retention rates with MNase coverage and CG methylation fractions obtained from [22] under Gene Expression Omnibus accessions “GSM1134620” and “GSM1134621”, respectively.

### Random Forest Modelling

We trained a Random Forest classifier to predict intron retention events using a comprehensive set of genomic and transcriptomic features. Because intron retention ratios are highly skewed and difficult to model accurately using regression, we framed the problem as a binary classification task. The classification target was defined as introns with retention ratios ≥0.025 (misspliced) versus <0.025 (properly spliced). Features included transcription levels (TPM), intron length, GC content, distance from the transcription start site, canonical splice site identity, frame disruption status, and one-hot encoded nucleotide sequences flanking the 5′ and 3′ splice sites (excluding the canonical dinucleotide positions). To prevent data leakage, we performed gene-level splitting, randomly partitioning genes into 80% training and 20% test sets such that all introns from a given gene were assigned to the same partition.

We optimized Random Forest hyperparameters (number of trees, maximum depth, minimum samples per split/leaf, and maximum features per split) using randomized search with 30 iterations and 3-fold grouped cross-validation on the training set, using ROC-AUC as the optimization metric. Class imbalance was addressed using balanced class weights. To identify a minimal feature set without sacrificing predictive performance, we implemented an automated feature selection procedure. We ranked features by their importance scores from the full model, then incrementally added in order of importance while tracking cumulative ROC-AUC on the held-out test set. We determined the optimal feature set using elbow detection via the Kneed algorithm. We produced partial dependence plots to visualize the marginal effect of each selected feature in the resulting minimal models on predicted intron retention probability. We also trained Random Forest models on autosomal introns using the same procedure. The extreme rarity of retention events in autosomal regions prevented effective model training, yielding low F1 scores (Table S4).

### M.pusilla Culture

The isolate of *M. pusilla* used here for RNA and chromatin analysis was obtained from the Roscoff Culture Collection. RCC834 is the same as CCMP1545, initially sequenced by Worden et al [7] and widely used as a reference for this species.

We grew cultures in low salinity (0.8X) artificial seawater with L1 supplements (except that Na_2_SiO_3_ was not added), [39] made using the L1 kit purchased from the National Center for Marine Algae/Bigelow (cat. No. MKL150L). We maintained small (10ml) cultures in vented T25 flasks at 18°C under a 14h light:10 h dark illumination cycle at 220 microEinsteins (uE).

To obtain cell pellets for RNA extraction, we transferred approximately 2 mL of the RCC834 culture into eight separate wells and maintained them in the above conditions for two days. On the third day, at 3 hours into the light cycle (1020 lux, 14 µE), we removed and pelleted each sample by centrifugation for 4 minutes at 5,000 g and 4°C, immediately flash froze the pellets in a dry ice-ethanol bath, and stored them at -80°C.

### Full-length Isoform Sequencing with R2C2 and Analysis with Mandalorion

We extracted RNA from approximately 1E8 cells of *M. pusilla* strain RCC834 (also known as CCMP1545) using a mixture of Trizol and chloroform, followed by purification with an RNeasy extraction kit. We prepared cDNA libraries through two sequential reactions. First, we performed reverse transcription on 200 ng of total RNA using Smartscribe reverse transcriptase, FS buffer, DTT, indexed oligo-dT primers, and a template switching oligo (TSO), incubating at 72°C for 3 minutes and 42°C for 60 minutes. Next, we amplified the resulting cDNA by PCR using KAPA HotStart master mix with primers targeting the oligo-dT and TSO sequences.

We circularized the cDNA libraries via Gibson assembly (NEBuilder HiFi) using a short, pre-annealed double-stranded DNA splint overlapping the cDNA ends. To remove uncircularized molecules, we digested the reaction with ExoI, ExoII, and Lambda Exonuclease (all NEB) for 16 hours at 37°C, then heat-inactivated the enzymes for 20 minutes at 80°C. We cleaned the reaction with SPRI beads at a 1:0.85 ratio.

We used the clean, circularized library as a template for rolling circle amplification (RCA) with Phi29 polymerase (NEB) and a random hexamer primer for 18 hours at 30°C, followed by heat inactivation for 10 minutes at 65°C. We then debranched the Phi29 reaction using T7 endonuclease for 2 hours at 37°C and cleaned and concentrated the product with a NEB Monarch DNA clean and concentrator column. We size-selected the resulting R2C2 library for DNA fragments larger than 3 kb using ProNex beads (Promega) and quantified DNA concentration with a Qubit fluorometer.

Table S6 lists the oligonucleotides used for library construction and circularization. We sequenced the multiplexed R2C2 library on an ONT PromethION R10.4 flow cell and basecalled reads with the Dorado basecaller using model dna_r10.4.1_e8.2_400bps_sup@v5.0.0.

The resulting raw reads were converted into consensus reads and demultiplexed using C3POa (v3.2). Based on these demultiplexed consensus reads, Mandalorion (v4.6) was used to identify and quantify high-confidence isoforms.

We refined the initial Mandalorion isoform calls using “sqanti3_qc” [40,41] using default parameters with the addition of “--isoAnnotLite” to attempt to liftover CDS features from the original gff3 file.

### Isoform Usage Analyses

We quantified isoform usage for each gene using Evenness, defined as the Shannon diversity index of transcript usage normalized by the maximum possible Shannon diversity for the observed number of isoforms. We fit linear models with log-transformed gene expression (log(TPM+1)) and intron number as covariates, and we included mating-type region status (inside vs. outside) as the main predictor. We tested the significance of each predictor, and the coefficient for mating-type region was highly significant, indicating that isoform usage is more even for genes inside the mating-type region after correcting for expression and intron number.

### Disruptions to protein function

We quantified possible effects on protein function in two ways. First, we compared the length of CDS annotated for each isoform with the CDS length in the canonical gene annotation. The expectation is that shorter CDS are less likely to be functional. Second, we first used interproscan (version 5.59-91.0) [42,43] with the flag “-goterms” to annotate functional domains and ontologies in all *M. pusilla* proteins. Then, to assess the effect of isoforms on protein function, we compared the proportion of isoforms that contained *X* percent of the total functional domains present for each gene across multiple values of *X*. We performed the same analysis for gene ontology terms. We then compared these results between genes in the MT region and the rest of the genome.

## Code Availability

Python scripts for processing data, performing statistical analyses, and plotting are available from the manuscript github at https://github.com/russcd/mating-type-missplicing. We also provide intermediate files containing summaries of the intron and isoforms for each species.

## Data Availability

Circularized consensus transcripts from the R2C2 and Mandalorion procedures are available under bioproject PRJNA1366634.

## Acknowledgements

The authors thank Alex Worden for insights about *Micromonas*, and Alexander Ioannidis for helpful comments on a previous version of this analysis and manuscript.

---

**Figure S1**. Intron length and GC content of introns in the autosomes (blue) and mating-type regions (orange) of Mamiellales species.

**Figure S2**. Sequence logos for each species at 5’ and 3’ splice sites for the autosomes, mating-type region and the delta between them. Data from M. pusilla is shown in Figure 2 and replicated here to facilitate comparison across species.

**Figure S3**. Nucleosome occupancy and methylation patterns in *M. pusilla* introns and flanking exons. (Top) MNase-based library preparation coverage within introns. (Middle) Average per CG dinucleotide modification rates in bisulfite sequencing data. (Bottom) Number of CG dinucleotide sites in the 50bp exons flanking each intron. We used a Mann-Whitney-U test to determine p-values. MNase and methylation data from [22].

**Figure S4:** Individual intron retention patterns across the chromosome containing the mating-type region of Mamiellales species. (Left), per intron retention frequency along autosomal regions (blue) and within the mating-type region (orange). (Right) Retention frequency for each intron plotted along each chromosome with colors as on the left panel. We obtained coordinates for mating type regions from [6]. Tick-marks indicate the midpoint of chromosome in each genome assembly.

**Table S1. Data sources and intron retention rates across Mamiellales species**.

**Table S2. Numbers of introns with a G at the 5th position in mating-type and non-mating type introns**.

**Table S3. Consensus branchpoint sequences were identified genome-wide using DREME motif discovery. ‘Present’ = exact 6/6 nucleotide match within the 50-nt branchpoint window upstream of the 3’ splice site. Ambiguity codes: V = A/C/G, R = A/G, S = C/G. Branchpoint is underlined**.

**Table S4. Random forest classifier model accuracy, F1 scores and feature importances**.

**Table S5. Protein structure preservation in annotated isoforms. Columns 2-4: proportion of isoforms retaining ≥X% of GO (Gene Ontology) terms assigned to the canonical protein. Columns 5-7: proportion of isoforms retaining ≥X% of annotated protein domains. These represent complementary measures of functional preservation**.

**Table S6. Oligonucleotides used in smartseq and R2C2 library construction**. Index corresponds to a random 8 nucleotide sequence unique to each replicate library.

**Table S7: NMD protein orthologs and their measured expression levels**.

