## Supplemental Table 7 for "Extensive splicing deficiency in a degenerating mating-type chromosome"

| **NMD Protein** | **M. pusilla ortholog** | **M. commoda ortholog** | **B. prasinos ortholog** | **O. tauri ortholog** | **M. pusilla TPM** | **M. commoda TPM** | **B. prasinos TPM** | **O. tauri TPM** |
| --- | --- | --- | --- | --- | --- | --- | --- | --- |
| UPF1 | MICPUCDRAFT_56002 | MICPUN_65270 | Bathy10g01700 | OT_ostta09g02800 | 28.0 | 55.2 | 48.5 | 39.7 |
| UPF2 | MICPUCDRAFT_46957 | MICPUN_58110 | Bathy18g00500 | OT_ostta04g01650 | 31.8 | 51.0 | 25.9 | 101.8 |
| UPF3 | NA | NA | NA | OT_ostta15g01265 | NA | NA | NA | 17.7 |
