## Supplemental Table 6 for "Extensive splicing deficiency in a degenerating mating-type chromosome"

| **Oligo ID** | **Sequence** |
| --- | --- |
| TSO-Smart-seq2 | AAGCAGTGGTATCAACGCAGAGTACATrGrGrG |
| Oligo_dT_Index1 | AAGCAGTGGTATCAACGCAGAGT [Index] ACTTTTTTTTTTTTTTTTTTTTTTTTTTTTTTVN |
| ISPCR primer | AAGCAGTGGTATCAACGCAGAGT |
| ISPCR_Anneal_Splint_1_F | ACTCTGCGTTGATACCACTGCTT  GCACGCACCGACAAACTCTGGCCGATTGTACTTTCTTATAAGGCGTAACACTAGACCATATCGTTTCTATAGATTAATTATGGCGAGAATTGCTAGCGAACTAGACAATTTTCGAAATAATCCTTTTTATAT AAGCAGTGGTATCAACGCAGAGT |
| ISPCR_Anneal_Splint_1_R | ACTCTGCGTTGATACCACTGCTT ATATAAAAAGGATTATTTCGAAAATTGTCTAGTTCGCTAGCAATTCTCGCCATAATTAATCTATAGAAACGATATGGTCTAGTGTTACGCCTTATAAGAAAGTACAATCGGCCAGAGTTTGTCGGTGCGTGC AAGCAGTGGTATCAACGCAGAGT |
