## Supplemental Table 5 for "Extensive splicing deficiency in a degenerating mating-type chromosome"

| **threshold (X)** | **mean proportion of transcripts containing X total functional ontology terms for MT genes** | **mean proportion of transcripts containing X total functional ontology terms for other genes** | **MWU P-value** | **mean proportion of transcripts containing X total functional domains for MT genes** | **mean proportion of transcripts containing X total functional domains for other genes** | **MWU P-value** |
| --- | --- | --- | --- | --- | --- | --- |
| 50 | 0.962 | 0.975 | 0.00454 | 0.950 | 0.968 | 0.0001041 |
| 60 | 0.957 | 0.972 | 0.000538 | 0.939 | 0.960 | 1.20E-05 |
| 70 | 0.944 | 0.967 | 9.22E-06 | 0.919 | 0.949 | 4.59E-08 |
| 80 | 0.930 | 0.961 | 8.06E-07 | 0.882 | 0.930 | 7.39E-10 |
| 90 | 0.928 | 0.958 | 6.78E-07 | 0.868 | 0.922 | 2.50E-10 |
