## Supplemental Table 4 for "Extensive splicing deficiency in a degenerating mating-type chromosome"

|  | Species | M. pusilla | M. commoda | B. prasinos | O. tauri |
| --- | --- | --- | --- | --- | --- |
| Evaluation | ROC-AUC (Full model) | 0.876 | 0.785 | 0.837 | 0.833 |
|  | ROC-AUC (Minimal model) | 0.867 | 0.735 | 0.812 | 0.830 |
|  | Mating-type F1 | 0.814 | 0.647 | 0.689 | 0.769 |
|  | Autosomal F1 | 0.061 | 0.000 | 0.167 | 0.000 |
| Feature Importances  (Minimal model) | TPM | 0.434 | 0.339 | 0.397 | 0.338 |
|  | Distance from TSS | 0.168 | 0.214 | 0.289 | 0.287 |
|  | Intron Length | 0.249 | 0.252 | 0.171 | 0.148 |
|  | Intron GC Content | 0.150 | 0.174 | 0.143 | 0.191 |
|  | Frame Disruption | Not Retained | 0.022 | Not Retained | Not Retained |
|  | C at position 1 of 3’ Exon | Not Retained | Not Retained | Not Retained | 0.036 |
