## Supplemental Table 3 for "Extensive splicing deficiency in a degenerating mating-type chromosome"

|  |  | Mating-Type | | | Non-mating type | | |  |
| --- | --- | --- | --- | --- | --- | --- | --- | --- |
|  | Branchpoint Sequence | Present | Absent | Fraction | Present | Absent | Fraction | P-value |
| M. pusilla | CTGACS | 109 | 914 | 0.107 | 3180 | 4787 | 0.399 | 3.64E-215 |
| M. commoda | CTVACC | 81 | 860 | 0.086 | 1741 | 2744 | 0.388 | 8.11E-85 |
| O. tauri | ACTRAC | 50 | 385 | 0.115 | 1219 | 392 | 0.757 | 3.58E-136 |
| B. prasinos | RCTSAC | 22 | 308 | 0.067 | 527 | 637 | 0.453 | 6.22E-45 |
