## Supplemental Table 2 for "Extensive splicing deficiency in a degenerating mating-type chromosome"

|  | Mating-Type | | | Non-mating type | | |  |
| --- | --- | --- | --- | --- | --- | --- | --- |
| Nucleotide in 5th intron position | G | Other | Fraction G | G | Other | Fraction G | Fisher’s Exact Test P-value |
| M. pusilla | 651 | 371 | 0.638 | 5668 | 2299 | 0.711 | 3.9E-05 |
| M. commoda | 617 | 324 | 0.656 | 3395 | 1090 | 0.757 | 1.6E-12 |
| O. tauri | 299 | 136 | 0.687 | 1475 | 136 | 0.916 | 1.4E-30 |
| B. prasinos | 155 | 175 | 0.470 | 812 | 352 | 0.698 | 3.7E-10 |
