## Supplemental Table 1 for "Extensive splicing deficiency in a degenerating mating-type chromosome"

| Species | Genome Accession | RNA-seq Accession | Mean Retention (Autosomes) | Mean Retention (Mating-Type) | Retention Mating-Type per Intron, per Transcript | Mean Retention Ratio (Mating-Type vs. Autosomal) |
| --- | --- | --- | --- | --- | --- | --- |
| M. pusilla | GCF_000151265.4 | SRR7814466 | 0.011 | 0.33 | 0.45 | 30 |
| M. commoda | GCF_000090985.2 | SRR7814480 | 0.0017 | 0.15 | 0.17 | 88 |
| B. prasinos | GCF_002220235.1 | SRR14396982 | 0.015 | 0.18 | 0.17 | 12 |
| O. tauri | GCF_000214015.3 | SRR5986291 | 0.010 | 0.2 | 0.23 | 20 |
