## Supplementary figures and images for "Extensive splicing deficiency in a degenerating mating-type chromosome"

### Supplemental Figure 1

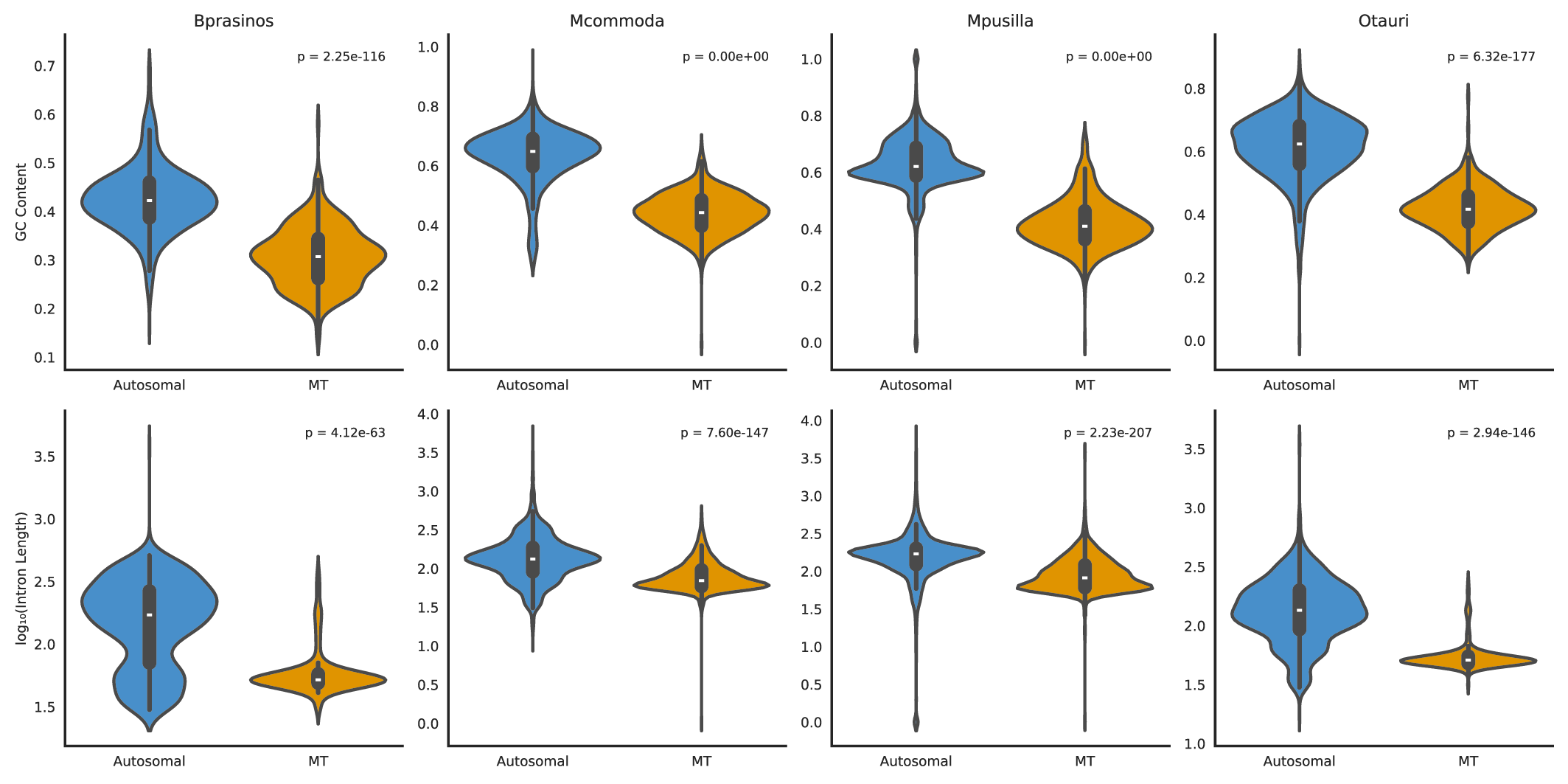

### Supplemental Figure 2

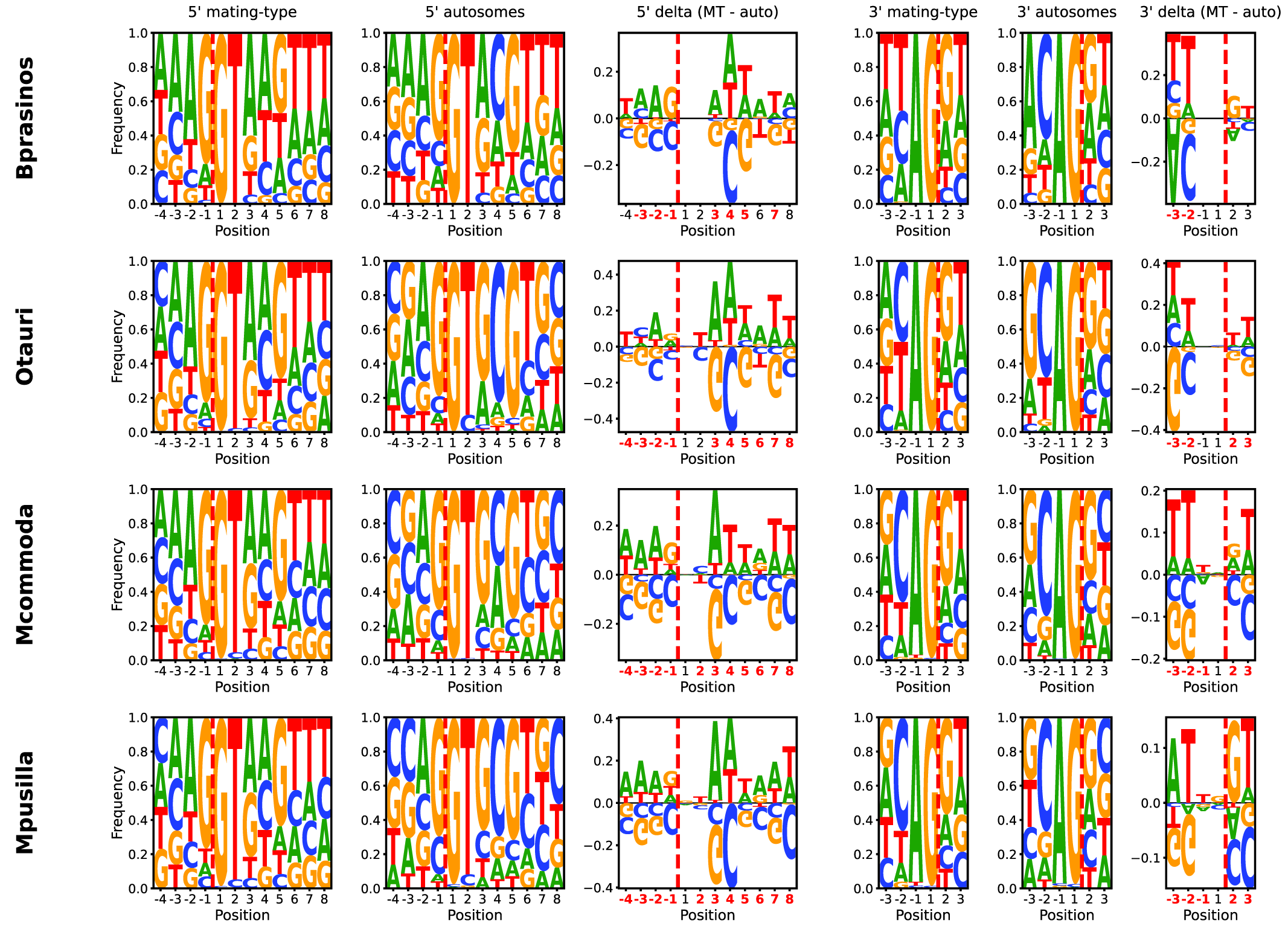

### Supplemental Figure 3

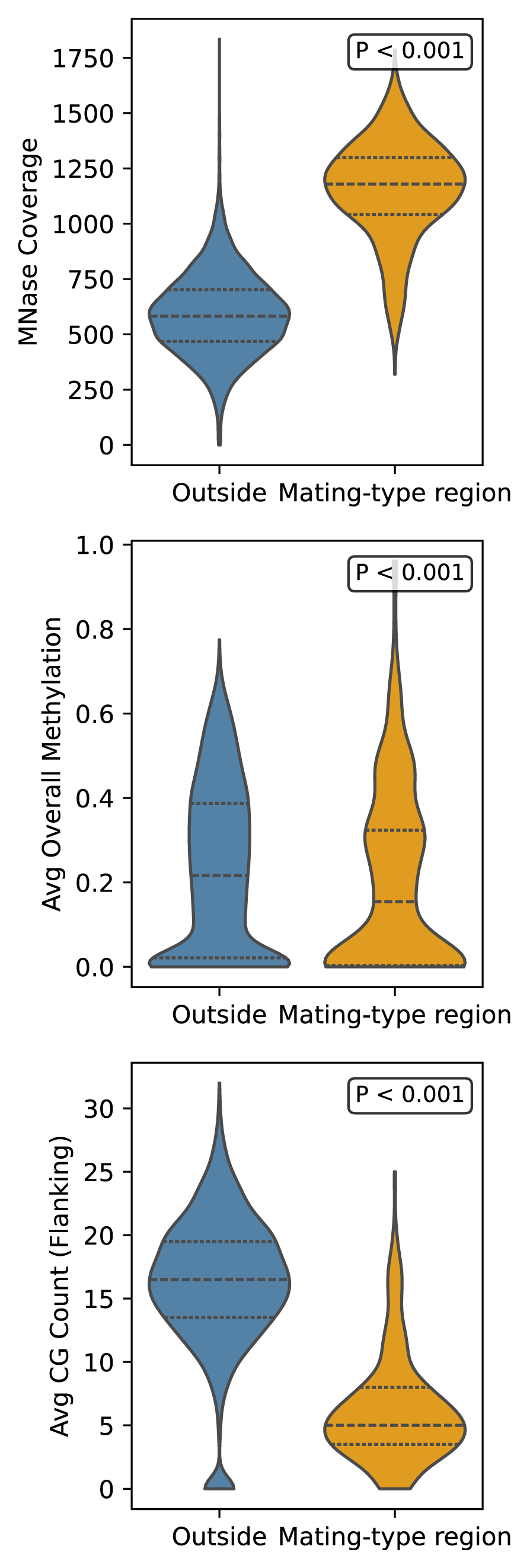

### Supplemental Figure 4

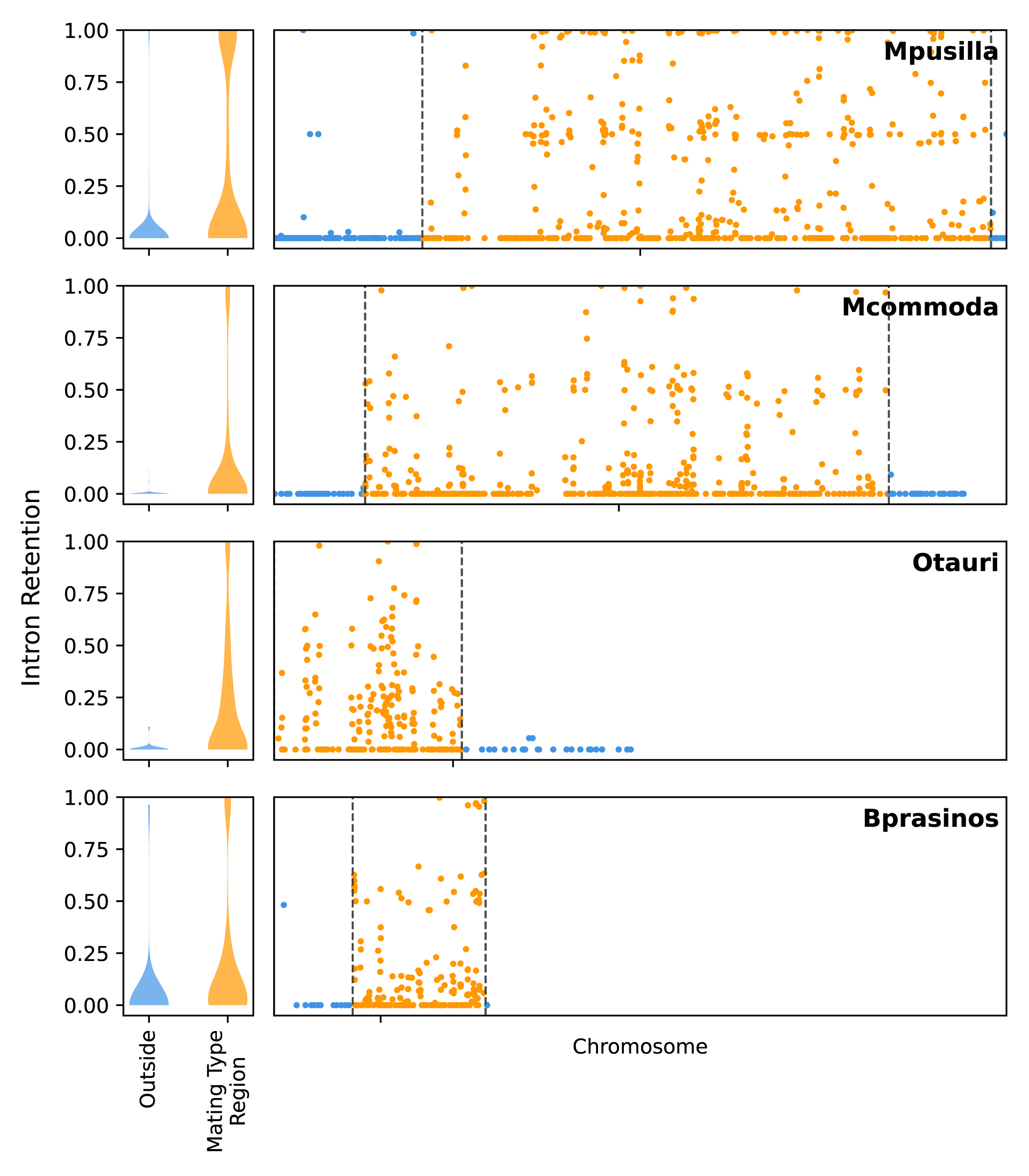
